# The renal hepcidin/ferroportin axis controls iron reabsorption and determines renal and hepatic susceptibility to iron overload

**DOI:** 10.1101/2020.07.02.183681

**Authors:** Goran Mohammad, Athena Matakidou, Peter A Robbins, Samira Lakhal-Littleton

## Abstract

The hepcidin/ferroportin axis controls systemic iron homeostasis by regulating iron acquisition from the duodenum and the reticuloendothelial system, respective sites of iron absorption and recycling. Ferroportin is also abundant in the kidney, where it has been implicated in iron reabsorption. However, it remains unknown whether hepcidin regulates ferroportin-mediated iron reabsorption and whether such regulation is important for systemic iron homeostasis. To address these questions, we generated a novel mouse model with an inducible renal-tubule specific knock-in of *fpnC326Y*, which encodes a hepcidin-resistant FPNC326Y. Under iron-replete conditions, female mice harbouring this allele had lower renal iron content and higher serum and liver iron levels than controls. Under conditions of excess iron availability, male and female mice harbouring this allele had greater liver iron overload, but lower renal iron overload relative to controls. In addition, hemochromatosis mice harbouring a ubiquitous knock-in of *fpnC326Y* did not develop renal iron overload otherwise seen in the setting of excess iron availability. These findings are the first formal demonstration that hepcidin regulates ferroportin-mediated iron reabsorption. They also show that loss of this regulation contributes to liver iron overload while protecting the kidney in the setting of hemochromatosis. Our findings have important implications. First, they indicate that targeting the hepcidin/ferroportin axis for treating iron overload disorders will inhibit iron reabsorption and increase renal iron content. Second, they suggest that inhibition of iron reabsorption by raised hepcidin in chronic inflammatory conditions contributes to iron deficiency and that parenteral iron supplementation in this setting may cause renal iron overload.

## INTRODUCTION

Ferroportin (FPN) is the only known mammalian iron export protein. It mediates iron release into the circulation from duodenal enterocytes and splenic reticuloendothelial macrophages, the respective sites of iron absorption and recycling (1,2). FPN-mediated iron release is antagonized by the hormone hepcidin, also known as hepcidin antimicrobial peptide (HAMP). Produced primarily in the liver, hepcidin binds to and induces internalization of FPN, thereby limiting iron release into the circulation and its availability to peripheral tissues (3,4). Thus, the HAMP/FPN axis operates at the sites of absorption and recycling to control systemic iron homeostasis.

FPN is also abundant in the kidney, the site of iron reabsorption (5–8). In this setting, it has been reported that FPN contributes to iron reabsorption by exporting filtered iron from renal tubules into the circulation (8). However, it remains unknown if renal FPN is also subject to regulation by HAMP, and if so, whether such regulation is important for systemic iron homeostasis.

To address these questions, we generated a novel mouse model with an inducible renal-tubule specific knock-in of *fpnC326Y*, which encodes a HAMP-resistant FPNC326Y protein. Additionally, to confirm the previously reported role of FPN in iron reabsorption, we also generated a mouse model with an inducible renal-tubule specific deletion of the *fpn* gene. We found that renal iron content was decreased by loss of HAMP responsiveness in renal tubules and increased by loss of FPN in renal tubules. Under iron replete conditions, these effects were confined to female mice and accompanied by transient changes in serum and liver iron levels. Under conditions of excess iron availability, loss of HAMP responsiveness in renal tubules exacerbated liver iron overload and reduced renal iron overload in male and female mice. Additionally, hemochromatosis mice harbouring a ubiquitous knock-in of *fpnC326Y* did not develop renal iron overload otherwise seen in iron-loaded wild type mice, despite similar degrees of liver iron overload in the two settings. Our findings are the first formal demonstration that HAMP directly controls FPN-dependent iron reabsorption. They show that this renal HAMP/FPN axis determines the degree of renal and hepatic iron overload in the setting of excess iron availability. They also demonstrate that loss of the renal HAMP/FPN axis contributes to liver iron overload while protecting the kidney in the setting of hemochromatosis.

Currently, there is considerable interest in strategies that target the HAMP/FPN axis for the treatment of diseases of iron overload (9). Our findings suggest that these strategies will additionally reduce iron reabsorption and increase renal iron content. Additionally, parenteral iron is increasingly being used to treat iron deficiency in chronic inflammatory conditions where serum HAMP is also raised, e.g. chronic kidney disease (10,11). Our findings suggest that inhibition of iron reabsorption may contribute to iron deficiency in these conditions and that increasing serum iron availability in this setting may cause renal iron overload.

## RESULTS

### The hepcidin/ferroportin axis in renal tubules controls iron reabsorption

To manipulate the FPN/HAMP axis in renal tubules, we used mice harbouring a Pax8.CreER^T2+^ knock-in transgene which drives tamoxifen-inducible expression of the Cre recombinase under control of the paired box gene 8 *pax8* promoter in proximal and distal tubules and in collecting ducts (12). To confirm further the efficacy and specificity of this Cre transgene, we crossed Pax8.CreER^T2+^ mice with mice harbouring the *Fpn*^fl/fl^ allele, and found that the presence of the deletion allele (ΔFpn) was indeed confined to the kidneys of *Fpn*^fl/fl^, Pax8.CreER^T2+^ mice (Supplemental Fig1 A).

To confirm the role of the FPN in renal iron homeostasis, we crossed Pax8.CreER^T2+^ mice with those harbouring a conditional knockout floxed *Fpn* allele. Furthermore, to understand the role of HAMP in renal iron homeostasis, we crossed Pax8.CreER^T2+^ mice with those harbouring a conditional knock-in floxed allele *FpnC326Y*, which encodes a hepcidin-resistant FPN. *Fpn*^fl/fl^,Pax8.CreER^T2+^ and *FpnC326Y*^fl/fl^,Pax8.CreER^T2+^ mice and their respective *Fpn*^fl/fl^ and *FpnC326Y*^fl/fl^ controls were induced with tamoxifen at 4 weeks of age, and their iron status characterised 1 week, 1 month, 3 months and 6 months later.

We found that renal iron content was higher in *Fpn*^fl/fl^,Pax8.CreER^T2+^ females than in the *Fpn*^fl/fl^ control females from 1 month onwards (Fig1A). Renal iron content in males was not different according to genotype (Fig1A). DAB-enhanced Perls iron stain confirmed iron accumulation within tubules of *Fpn*^fl/fl^,Pax8.CreER^T2+^ females (Fig1B). Conversely, renal iron content was lower in *FpnC326Y*^fl/fl^,Pax8.CreER^T2+^ females than in *FpnC326Y*^fl/fl^ control females from 1 month onwards (Fig1C). Renal iron content in males was not different according to genotype (Fig1C). DAB-enhanced Perls iron stain also confirmed reduction in iron levels within tubules of *FpnC326Y*^fl/fl^,Pax8.CreER^T2+^ females (Fig1D). Serum iron levels were decreased in *Fpn*^fl/fl^,Pax8.CreER^T2+^ females relative to *Fpn*^fl/fl^ control females at 1 and 3 month timepoints, while they remained comparable in males of different genotypes at all timepoints (Fig1E). Conversely, serum iron levels were increased transiently both in male and in female *FpnC326Y*^fl/fl^,Pax8.CreER^T2+^ mice relative to *FpnC326Y*^fl/fl^ controls at the 1 month timepoint (Fig1F). Increased renal iron content and decreased serum iron levels in *Fpn*^fl/fl^,Pax8.CreER^T2+^ females confirm that renal FPN contributes to iron reabsorption. Additionally, decreased renal content and increased serum iron levels in *FpnC326Y*^fl/fl^,Pax8.CreER^T2+^ females demonstrate that FPN-dependent iron reabsorption is subject to regulation by HAMP. These data also indicate that, under normal physiological conditions, the control of iron reabsorption by the renal HAMP/FPN axis is more important in females than in males.

**Figure 1.**
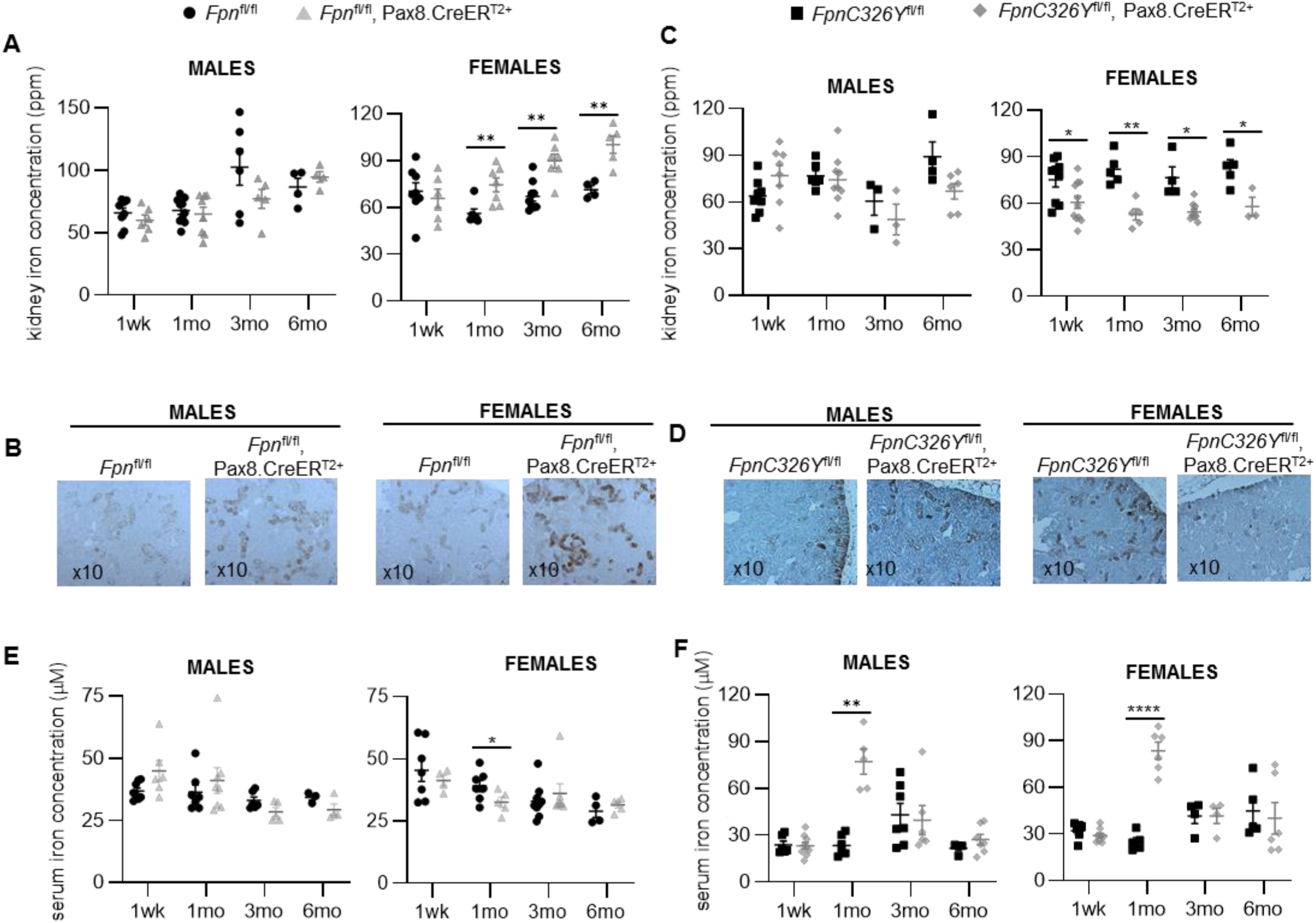
The HAMP/FPN axis in renal tubules controls iron-reabsorption. **A.** Renal iron levels in male and female *Fpn*^fl/fl^,Pax8.CreER^T2+^ mice and *Fpn*^fl/fl^ controls at 1 week, 1 month, 3 months and 6 months post-tamoxifen induction. **B.** Representative images of DAB-enhanced Perls iron stain in kidneys of corresponding animals at 3 months post tamoxifen induction. **C** Renal iron levels in male and female *FpnC326Y*^fl/fl^,Pax8.CreER^T2+^ mice and *FpnC326Y*^fl/fl^ controls at 1 week, 1 month, 3 months and 6 months post-tamoxifen induction. **D.** Representative images of DAB-enhanced Perls iron stain in kidneys of corresponding animals at 3 months post tamoxifen induction. **E.** Serum iron levels in male and female *Fpn*^fl/fl^,Pax8.CreER^T2+^ mice and *Fpn*^fl/fl^ controls at 1 week, 1 month, 3 months and 6 months post-tamoxifen induction. **F.** Serum iron levels in male and female *FpnC326Y*^fl/fl^,Pax8.CreER^T2+^ mice and *FpnC326Y*^fl/fl^ controls at 1 week, 1 month, 3 months and 6 months post-tamoxifen induction. Values are shown as mean± standard error of the mean. *p<0.05, **p<0.01, ****p<0.0001.

### The renal HAMP/FPN axis contributes normal systemic iron homeostasis under iron-replete conditions

Next, we set out to determine the contribution of the renal HAMP/FPN axis to systemic iron homeostasis. We found that *Fpn*^fl/fl^,Pax8.CreER^T2+^ females have lower serum ferritin levels at the 1 and 3 month timepoints (Fig2A), and lower liver iron content at the 3 and 6 month timepoints, when compared to *Fpn*^fl/fl^ control females (Fig2B). They also had a transient reduction in haemoglobin levels at the 3 month timepoint (Fig2C). None of these parameters were different between *Fpn*^fl/fl^,Pax8.CreER^T2+^ males and their *Fpn*^fl/fl^ controls (Fig2A-C). In contrast, *FpnC326Y*^fl/fl^,Pax8.CreER^T2+^ females had a transient rise in serum ferritin (Fig2D) and in liver iron content (Fig2E) at the 1 month timepoint when compared to *FpnC326Y*^fl/fl^ control females. Their haemoglobin levels remained comparable to those of controls at all timepoints (Fig2F). None of these parameters were different between *FpnC326Y*^fl/fl^,Pax8.CreER^T2+^ males and their *FpnC326Y*^fl/fl^ controls (Fig2D-F). Together, these data demonstrate that, under normal physiological conditions, the renal HAMP/FPN axis contributes to systemic iron homeostasis but that this contribution is minor compared to that of the HAMP/FPN axes in the duodenum and reticuloendothelial system.

**Figure 2.**
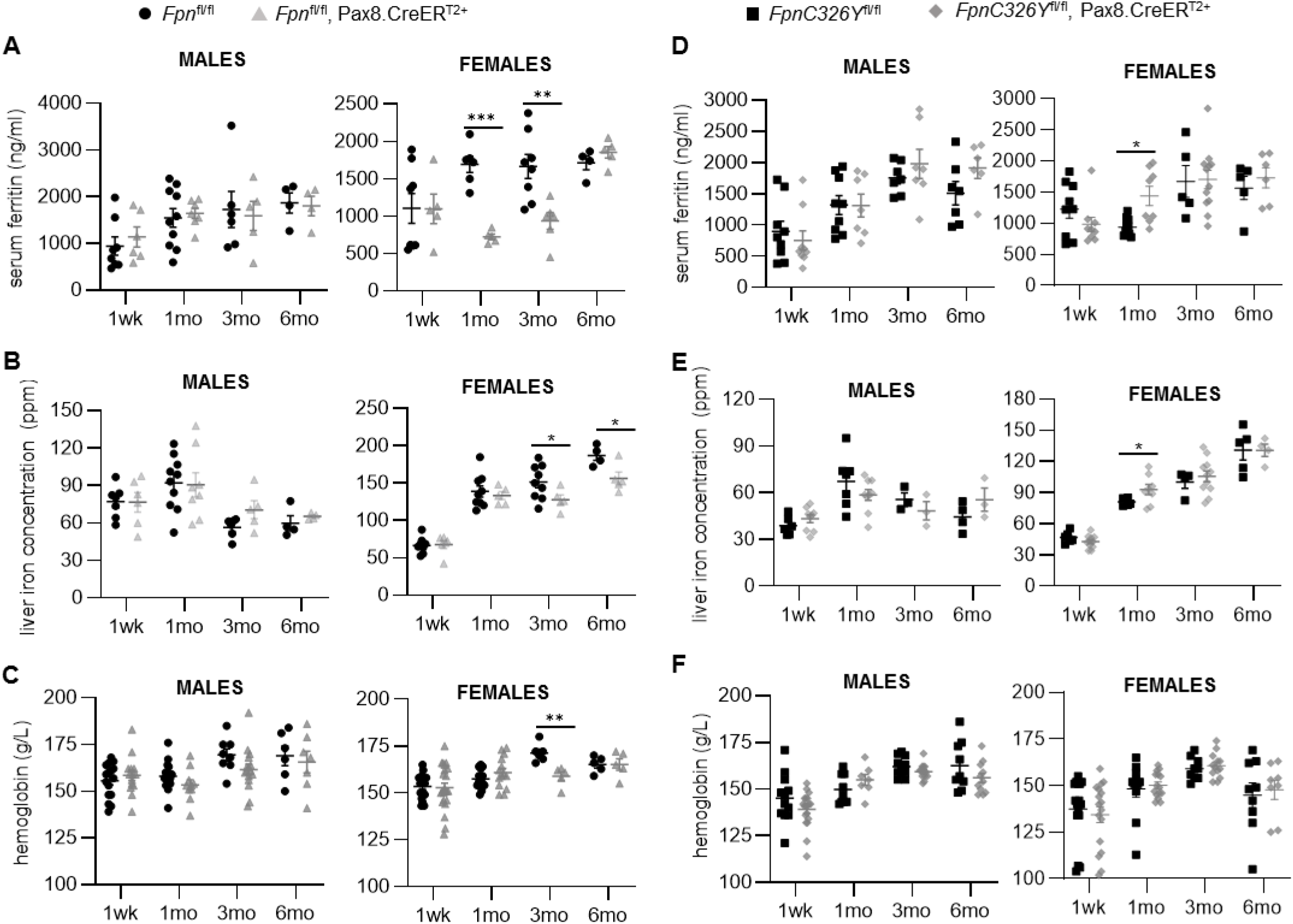
The renal HAMP/FPN axis contributes to systemic iron homeostasis. **A.** Serum ferritin levels in male and female *Fpn*^fl/fl^,Pax8.CreER^T2+^ mice and *Fpn*^fl/fl^ controls at 1 week, 1 month, 3 months and 6 months post-tamoxifen induction. **B.** Liver iron concentration in male and female *Fpn*^fl/fl^,Pax8.CreER^T2+^ mice and *Fpn*^fl/fl^ controls at 1 week, 1 month, 3 months and 6 months post-tamoxifen induction. **C.** Hemoglobin levels in male and female *Fpn*^fl/fl^,Pax8.CreER^T2+^ mice and *Fpn*^fl/fl^ controls at 1 week, 1 month, 3 months and 6 months post-tamoxifen induction. **D.** Serum ferritin levels in male and female *FpnC326Y*^fl/fl^,Pax8.CreER^T2+^ mice and *FpnC326Y*^fl/fl^ controls at 1 week, 1 month, 3 months and 6 months post-tamoxifen induction. **E.** Liver iron concentration in male and female *FpnC326Y*^fl/fl^,Pax8.CreER^T2+^ mice and *FpnC326Y*^fl/fl^ controls at 1 week, 1 month, 3 months and 6 months post-tamoxifen induction. **F.** Hemoglobin levels in male and female *FpnC326Y*^fl/fl^,Pax8.CreER^T2+^ mice and *FpnC326Y*^fl/fl^ controls at 1 week, 1 month, 3 months and 6 months post-tamoxifen induction. Values are shown as mean± standard error of the mean. *p<0.05, **p<0.01, ***p<0.001.

### The renal HAMP/FPN axis determines the degree of renal and hepatic iron loading under conditions of excess iron availability

Next, we set out to determine the role of the renal HAMP/FPN axis in the setting of iron overload. To that effect, *FpnC326Y*^fl/fl^,Pax8.CreER^T2+^ animals and *FpnC326Y*^fl/fl^ controls were induced with tamoxifen then provided either a control chow diet containing 200ppm iron, or an iron-loaded diet containing 5000ppm iron for 3 months. We found that provision of iron-loaded diet increased renal and liver iron content in males and females of all genotypes. However, the degree of renal iron loading was lower in *FpnC326Y*^fl/fl^,Pax8.CreER^T2+^ animals than in *FpnC326Y*^fl/fl^ controls (Fig3A). This result was confirmed by DAB-enhanced Perls iron stain of kidneys (Fig3B). In contrast, the degree of liver iron loading was higher in *FpnC326Y*^fl/fl^,Pax8.CreER^T2+^ animals than in *FpnC326Y*^fl/fl^ controls (Fig3C) and this was also confirmed by DAB-enhanced Perls stain (Fig3D). These findings demonstrate that, under conditions of excess iron availability, the control of FPN-dependent iron reabsorption by HAMP decreases liver iron overload while increasing renal iron overload (Fig5A).

**Figure 3.**
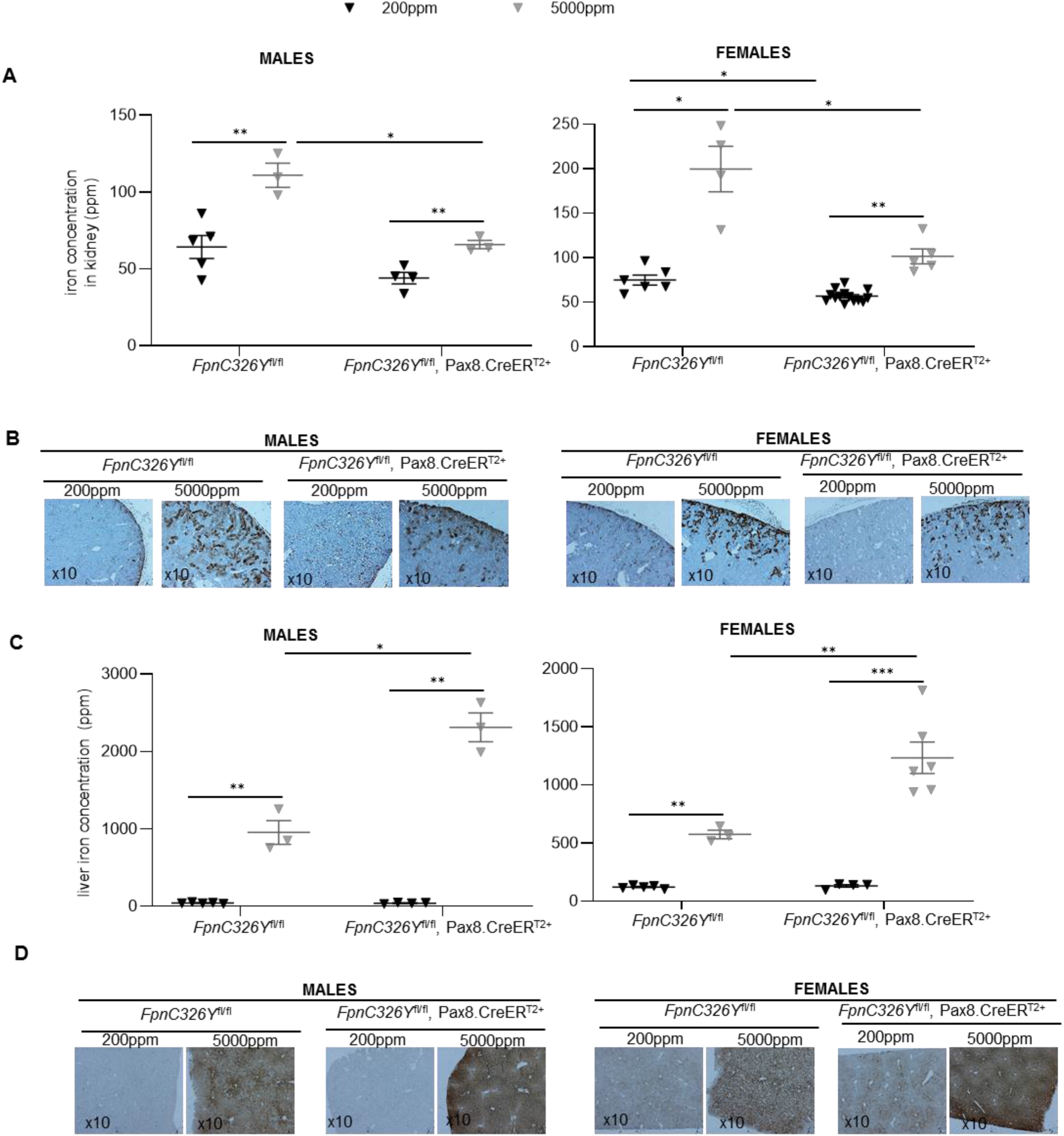
The renal HAMP/FPN axis determines the degree or renal and hepatic iron loading under conditions of excess iron availability. **A.** Renal iron levels in male and female *FpnC326Y*^fl/fl^,Pax8.CreER^T2+^ mice and *FpnC326Y*^fl/fl^ controls, provided either a control chow diet (200ppm) or an iron-loaded diet (5000ppm) for 3 months. **B.** Representative images of DAB-enhanced Perls iron stain in kidneys of corresponding animals. **C**. Liver iron levels in male and female *FpnC326Y*^fl/fl^,Pax8.CreER^T2+^ mice and *FpnC326Y*^fl/fl^ controls, provided either a control chow diet (200ppm) or an iron-loaded diet (5000ppm) for 3 months. **D.** Representative images of DAB-enhanced Perls iron stain in kidneys of corresponding animals. Values are shown as mean± standard error of the mean. *p<0.05, **p<0.01, ***p<0.001.

### The renal HAMP/FPN axis determines the pattern of tissue iron overload in hemochromatosis

Next, we set out to explore the role of the renal HAMP/FPN axis in the context of hereditary hemochromatosis, a genetic condition of iron overload caused by defects in hepcidin production or hepcidin responsiveness (13). To that effect, we used mice generated in-house harbouring a heterozygous ubiquitous knock-in of the fpnC326Y allele (*Fpn*^wt/C326Y^). We had previously demonstrated that these mice develop the iron-overload phenotype characteristic of hereditary hemochromatosis (14, 15). We found that while *Fpn*^wt/C326Y^mice had higher liver iron content than their *Fpn*^wt/wt^ controls, their renal iron content was normal (Fig4A, B). In contrast, wild type animals provided an iron-loaded diet (5000ppm iron for 3 months) developed both liver and renal iron overload (Fig4C,D). FPN levels in renal tubules were raised both in *Fpn*^wt/C326Y^mice and in wild type mice provided an iron-loaded diet (Fig4E,F). However, FPN localised primarily to the cytoplasm and basolateral membrane in renal tubules of *Fpn*^wt/C326Y^mice, while it localised primarily to cytoplasm and the apical membrane in wild type mice provided an iron-loaded diet (Fig4E,F). These observations, together with the previous finding that *FpnC326Y*^fl/fl^,Pax8.CreER^T2+^ animals have lower renal iron loading than controls following provision of an iron-loaded diet, demonstrate that loss of HAMP action on renal FPN protects the kidney from iron loading in the setting of hereditary hemochromatosis (Fig5A,B).

**Figure 4.**
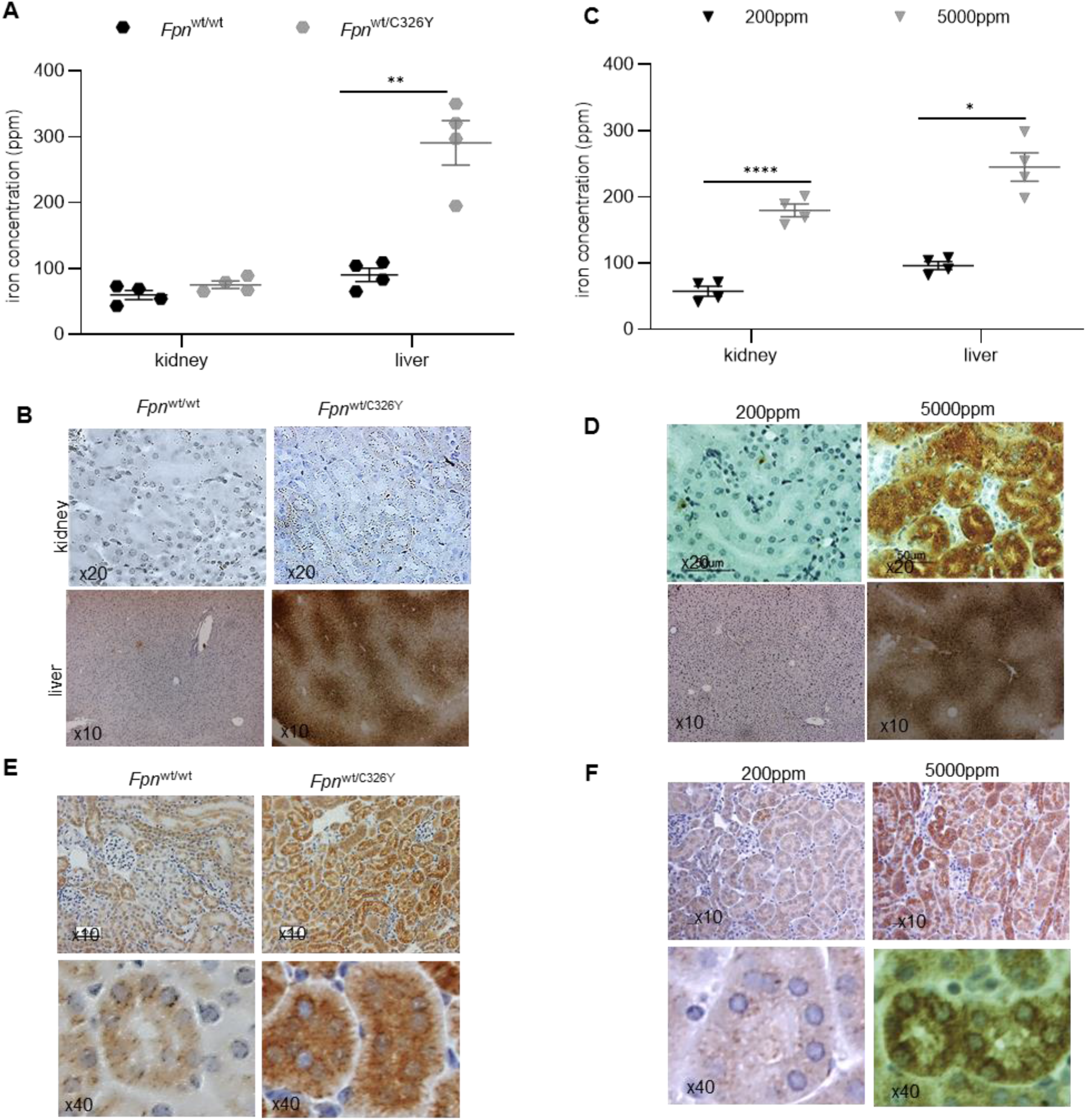
The renal HAMP/FPN determines the pattern of tissue iron overload in hemochromatosis. **A.** Renal and hepatic iron levels in *Fpn*^wt/C326Y^ animals and *Fpn*^wt/wt^ controls at 3 months of age. **B.** Representative images of DAB-enhanced Perls iron stain in kidneys and livers of corresponding animals. **C.** Renal and hepatic iron levels in wild type animals provided either a control chow diet (200ppm) or an iron-loaded diet (5000ppm) from weaning for 3 months. **D.** Representative images of DAB-enhanced Perls iron stain in kidneys and livers of corresponding animals. **E.** Representative images of FPN immunostaining in kidneys of *Fpn*^wt/C326Y^ animals and *Fpn*^wt/wt^ controls at 3 months of age. **F.** Representative images of FPN immunostaining in kidneys of wild type animals provided either a control chow diet (200ppm) or an iron-loaded diet (5000ppm) from weaning for 3 months. Values are shown as mean± standard error of the mean. *p<0.05, **p<0.01, ****p<0.0001.

**Figure 5.**
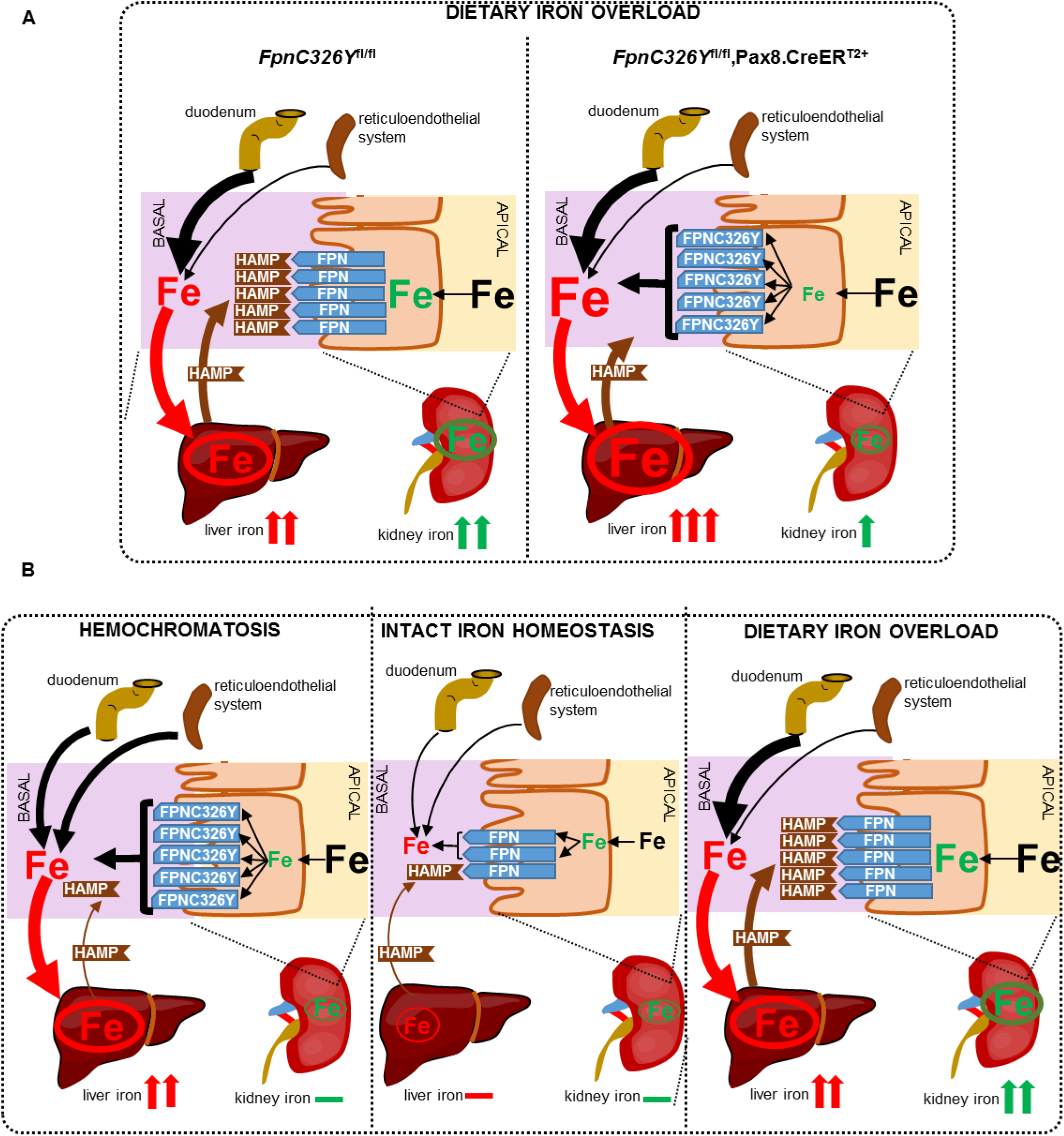
The role of the renal HAMP/FPN axis in the setting of iron overload. **A.** Provision of iron-loaded diet increases serum iron availability and iron levels in the glomerular filtrate and raises hepcidin (HAMP) production and release by the liver. Raised serum HAMP inhibits iron-reabsorption by blocking FPN in renal tubules. Inhibition or iron reabsorption causes renal iron retention while decreasing systemic iron availability and consequently reducing liver iron overload. In *FpnC3326Y*^fl/fl,^ Pax8.CreER^T2+^ mice, loss of HAMP responsiveness in the renal tubules causes unregulated iron reabsorption. This in turn prevents renal iron retention while increasing systemic iron availability and subsequently increasing liver iron overload. **B.** In mice with intact iron homeostasis, provision of iron-loaded diet increases serum iron levels and iron levels in the glomerular filtrate. However, iron reabsorption is blocked by HAMP which is elevated in this setting. The combination of increased iron in the glomerular filtrate and decreased iron reabsorption leads to renal iron overload. In contract, iron reabsorption from renal tubules is enhanced in hemochromatosis mice (Fpn^wt/C326Y^), due to loss of HAMP responsiveness in renal tubules. Enhanced iron reabsorption mitigates against the effects of increased iron levels in the glomerular filtrate, protecting the kidney from iron overload.

## DISCUSSION

The major finding of the present study is that HAMP regulates iron reabsorption by controlling FPN in renal tubules. Loss of this regulation through knock-in of HAMP-resistant FPN in renal tubules decreased renal iron content and increased serum iron and liver iron stores. Previous studies had reported the regulation of FPN by HAMP in cultured renal cells, and an inverse relationship between HAMP and FPN levels in the kidney following unilateral ureter occlusion (6, 7). The present study provides the first in-vivo demonstration of a direct role for HAMP in the regulation of iron reabsorption, and of the importance of this regulation in the context of systemic iron homeostasis.

The second important finding of the present study is that the renal HAMP/FPN axis is physiologically important in the setting of excess iron availability because it determines the magnitude of renal and hepatic iron overload. Indeed, loss of HAMP responsiveness in renal tubules exacerbated the magnitude of liver iron overload while reducing the magnitude of renal iron overload following provision of an iron-loaded diet. This finding indicates that unregulated iron reabsorption contributes to liver iron overload in the setting of hemochromatosis. Additionally, renal iron overload was observed in the setting of excess dietary iron (wild type mice in whom the renal HAMP/FPN axis in intact) but not in the setting of genetic iron overload (mice with ubiquitous knock-in of *fpnC326Y*, in whom the renal HAMP/FPN axis is also disrupted). This finding explains the clinical observation that the kidney is not commonly affected in patients with hereditary hemochromatosis (13).

Interestingly, FPN levels in renal tubules were enhanced both in the settings of hemochromatosis and of dietary iron overload. In the setting of hemochromatosis, enhanced FPN levels in the renal tubules in the absence of any intracellular iron deposition, and the localisation of FPN to the basolateral membrane likely reflect the result of loss of FPN regulation by serum HAMP. In contrast, enhanced FPN levels in the setting of dietary iron overload likely reflect an IRP-driven response to increased intracellular iron levels within renal tubules, while its cytoplasmic/apical localisation likely reflect post-translational inhibition by serum HAMP (which is raised in response to dietary iron loading). As such, FPN localisation within renal tubules is determined both by the concentration of HAMP in the serum and by the levels of filtered iron in renal tubules. This finding may explain the divergence between previous reports as to the localisation of FPN in renal tubules (5–8,16, 17). The functional significance of the apical localisation of FPN in renal tubules is not known. In the future, it would be important to determine whether the presence of FPN on the apical membrane is involved in iron excretion into the urine, and if so, whether such excretion is subject to regulation by HAMP in the tubular lumen. Another unanswered question is the extent to which HAMP derived from the renal tissue itself controls FPN. Relevant to this, we found that expression of the *hamp* gene in renal tubules was raised following provision of iron-loaded diet to wild type mice but not in hemochromatosis mice, suggesting that renal HAMP expression may be regulated by intracellular iron levels within tubules (Supplemental Fig2). In the future, it would be interesting to determine the relative contributions of renal and hepatic HAMPs to the control of iron reabsorption.

Another important finding of the present study is that, under normal physiological conditions, the renal HAMP/FPN axis contributes to systemic iron homeostasis but that this contribution is minor compared to that of the HAMP/FPN axes in the duodenum and reticuloendothelial system. Indeed, the observed changes in serum and liver iron indices resulting from loss of FPN or of HAMP-responsiveness in the real tubules were transient, suggesting the involvement compensatory mechanism(s). One possible compensatory mechanism is modulation of hepatic HAMP in response to changes in serum iron availability. Consistent with this notion, hepatic *hamp* gene expression was decreased in livers of female mice with renal-tubule specific loss of FPN and increased in livers of female mice with renal-tubule specific loss of HAMP-responsiveness (Supplemental Fig3). As well as being transient in nature, the observed changes resulting from loss of FPN or of HAMP-responsiveness in the renal tubules were confined to female mice. This finding could not be attributed to differences between males and females in the activity of the Pax8.CreERT^2+^ transgene because the product of fpn knockout (ΔFpn) was detected in kidneys of both male and female *fpn*^fl/fl^, Pax8.CreERT^2+^ mice (Supplemental Fig1A). Instead, this finding suggests that, at least in C57BL/6 mice, the contribution of renal iron reabsorption to systemic iron levels is more important in females than in males. A previous study using a different Cre recombinase transgene, driven by a constitutively active Nestin promoter to delete *fpn* in entire nephron also reported an increase in renal iron levels, and decrease in serum iron and liver iron stores, although that study was conducted in a different mouse strain (129/SvEvTac), did not distinguish between males and females and did not report on the timecourse of these changes (8).

The findings of the present study have potentially important clinical implications. First, they suggest that strategies targeting the HAMP/FPN axis for the treatment of iron overload e.g. HAMP mimetics, FPN inhibitors, may reduce iron reabsorption and increase renal iron content (9). Second, they indicate that inhibition of iron reabsorption by HAMP may contribute to iron deficiency in the setting of chronic conditions, where HAMP is raised by inflammation, e.g. chronic kidney disease. Finally, they suggest that raising systemic iron availability in this setting (e.g. using parenteral iron supplementation) may affect renal function by causing renal iron overload (10,11).

## METHODS

### Mice

All animal procedures were compliant with the UK Home Office Animals (Scientific Procedures) Act 1986 and approved by the University of Oxford Medical Sciences Division Ethical Review Committee.

The conditional fpn^fl^, and fpnC326Y^fl^ alleles was generated as described previously (14, 18). Mice harbouring the Pax8.CreERT2+ transgene were a gift from Dr Athena Matakidou, Cancer Research UK Cambridge Institute, University of Cambridge. These mice were generated as described previously (12).

### Diets

Unless otherwise stated, animals were provided with a standard rodent chow diet containing 200ppm iron. In iron manipulation experiments, mice were given an iron-loaded diet (5,000ppm iron; Teklad TD.140464) or a matched control diet (200ppm iron; Teklad TD.08713) from weaning for 3 months.

### Iron quantitation

Serum iron and ferritin levels were determined using the ABX-Pentra system (Horiba Medical, CA). Determination of total elemental iron in tissues was carried out by inductively coupled plasma mass spectrometry (ICP-MS) as described previously (14, 15, 18). Calibration was achieved using the process of standard additions, where spikes of 0ng/g, 0,5ng/g, 1ng/g, 10ng/g, 20ng/g and 100ng/g iron were added to replicates of a selected sample. An external iron standard (High Purity Standards ICP-MS-68-A solution) was diluted and measured to confirm the validity of the calibration. Rhodium was also spiked onto each blank, standard and sample as an internal standard at a concentration of 1ng/g. Concentrations from ICP-MS were normalised to starting tissue weight.

### Immunohistochemistry

Formalin-fixed paraffin-embedded tissue sections were stained with rabbit polyclonal anti-mouse FPN antibody (MTP11-A, Alpha Diagnostics, RRID:AB_1619475) at 1/200 dilution.

### DAB-enhanced Perls stain

Formalin-fixed paraffin-embedded tissue sections were deparaffinised using Xylene, then rehydrated in ethanol. Slides were then stained for 1 hour with 1% potassium ferricyanide in 0.1mol/L HCl buffer. Endogenous peroxidase activity was quenched, then slides were stained with DAB chromogen substrate and counterstained with haematoxylin. They were visualised using a standard brightfield microscope.

### Quantitative PCR

Gene expression was measured using Applied Biosystems Taqman gene expression assay probes for *Hamp* and house-keeping gene *β-Actin* (Life Technologies, *Carlsbad, CA*). The CT value for the gene of interest was first normalised by deducting CT value for *β-Actin* to obtain a delta CT value. Delta CT values of test samples were further normalised to the average of the delta CT values for control samples to obtain delta delta CT values. Relative gene expression levels were then calculated as 2^−delta deltaCT^.

### Statistics

Values are shown as mean± standard error of the mean (S.E.M). Paired comparisons were performed using Student’s T test. Multiple comparisons were drawn using ANOVA. Post-hoc tests used Bonferroni correction.

## ACKNOWLEDGMENTS

S L-L was the recipient of a British Heart Foundation Intermediate Basic Science Postdoctoral Fellowship (FS/12/63/29895). GM was funded by a Kidney Research UK project grant (RP_020_20160303) awarded to S L-L.

---

**Supplemental Figure 1.**
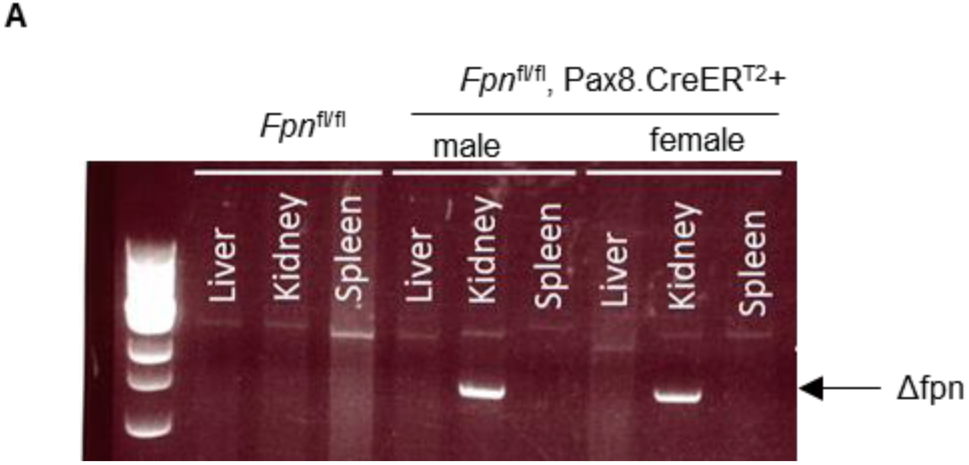
Confirmation of Pax8.CreER^T2+^ activity and specificity. Genotyping of *Fpn*^fl/fl^ and *Fpn*^fl/fl^,Pax8.CreER^T2+^ mice using genomic DNA extracted from the liver, spleen and kidney at 1 week post tamoxifen induction.

**Supplemental Figure 2.**
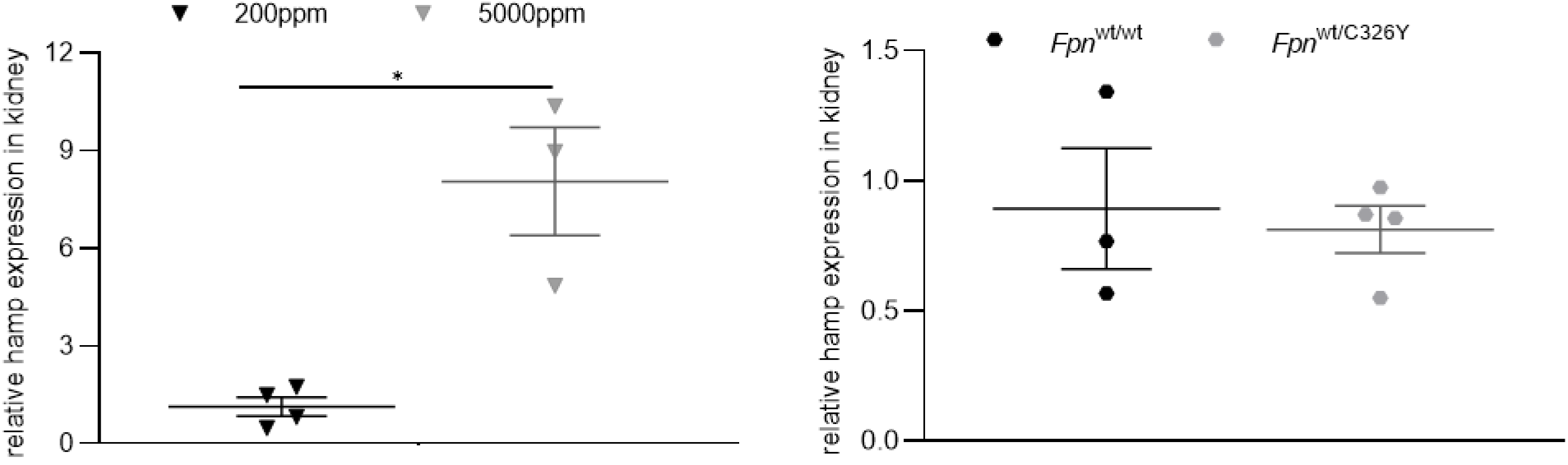
Renal hamp expression is increased by dietary iron overload but not in hemochromatosis. **A.** Relative expression of hamp mRNA in kidneys of *Fpn*^wt/C326Y^ animals and *Fpn*^wt/wt^ controls at 3 months of age. **B.** Relative expression of hamp mRNA in kidneys of wild type animals fed an iron-loaded diet containing 5000ppm iron, or a control diet containing 200ppm iron from weaning for 3 months. Values are shown as mean± standard error of the mean. *p<0.05.

**Supplemental figure 3.**
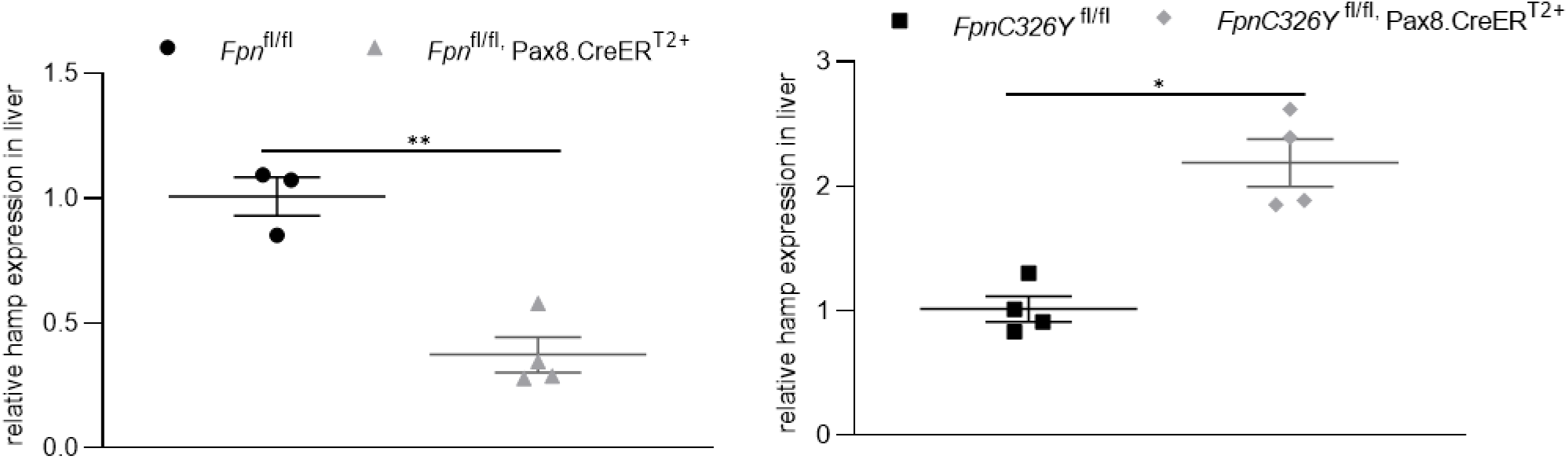
Liver hamp expression is decreased in mice with renal tubule-specific knockout of *fpn* and increased in mice with renal tubule-specific knock-in of *fpnC326Y*. **A.** Relative expression of hamp mRNA in livers of female *Fpn*^fl/fl,^ Pax8.CreER^T2+^ mice and *Fpn*^fl/fl^ controls at 1 month timepoint. **B.** Relative expression of hamp mRNA in livers of female *FpnC326Y*^fl/fl,^ Pax8.CreER^T2+^ mice and *FpnC326Y*^fl/fl^ controls at 1 month timepoint. Values are shown as mean± standard error of the mean. *p<0.05, **p<0.01.

## REFERENCES

1. Donovan A, Lima CA, Pinkus JL, Pinkus GS, Zon LI, Robine S, Andrews NC. The iron exporter ferroportin/slc40a1 is essential for iron homeostasis. Cell metabolism. 2005;1:191–200

2. Ganz T. Cellular iron: Ferroportin is the only way out. Cell metabolism. 2005;1:155–157

3. Nemeth E, Tuttle MS, Powelson J, Vaughn MB, Donovan A, Ward DM, Ganz T, Kaplan J. Hepcidin regulates cellular iron efflux by binding to ferroportin and inducing its internalization. Science. 2004;306:2090–2093

4. Qiao B, Sugianto P, Fung E, Del-Castillo-Rueda A, Moran-Jimenez MJ, Ganz T, Nemeth E. Hepcidin-induced endocytosis of ferroportin is dependent on ferroportin ubiquitination. Cell metabolism. 2012;15:918–924

5. Wolff N, Liu W, Fenton R, Lee W, Thevenod F, Smith C. Ferroportin 1 is expressed basolaterally in rat kidney proximal tubule cells and iron excess increases its membrane trafficking. Journal of cellular and molecular medicine. 2011;15:209–219.

6. Pan S, Qian ZM, Cui S, et al. Local hepcidin increased intracellular iron overload via the degradation of ferroportin in the kidney. Biochem Biophys Res Commun. 2020;522(2):322–327.

7. Moulouel B, Houamel D, Delaby c, et al. Hepcidin regulates intrarenal iron handling at the distal nephron. Kidney International. 2013;84:756–766

8. Wang X, Zheng X, Zhang J, et al. Physiological functions of ferroportin in the regulation of renal iron recycling and ischemic acute kidney injury. AJP: Renal Physiology. 2018;315(4):F1042‐F1057.

9. Katsarou A, Pantopoulos K. Hepcidin Therapeutics. Pharmaceuticals (Basel). 2018;11(4):127.

10. Charytan C, Bernardo MV, Koch TA, Butcher A, Morris D, Bregman DB. Intravenous ferric carboxymaltose versus standard medical care in the treatment of iron deficiency anemia in patients with chronic kidney disease: a randomized, active-controlled, multi-center study. Nephrology Dialysis and Transplant. 2013;28:953–964. 12.

11. Macdougall IC, Bock AH, Carrera F, Eckardt KU, Gaillard C, Van Wyck D, Roubert B, Nolen JG, Roger SD, FIND-CKD Study Investigators FIND-CKD: a randomized trial of intravenous ferric carboxymaltose versus oral iron in patients with chronic kidney disease and iron deficiency anaemia. Nephrology, Dialysis and Transplant. 2014;29:2075–2084

12. Espana-Agusti J, Zou X, Wong K, et al. Generation and Characterisation of a Pax8-CreERT2 Transgenic Line and a Slc22a6-CreERT2 Knock-In Line for Inducible and Specific Genetic Manipulation of Renal Tubular Epithelial Cells. PLoS One. 2016;11(2):e0148055.

13. Camaschella C. Understanding iron homeostasis through genetic analysis of hemochromatosis and related disorders. Blood. 2005;106:3710–3717

14. Lakhal-Littleton S, Wolna M, Chung YJ, et al. An essential cell-autonomous role for hepcidin in cardiac iron homeostasis. Elife. 2016;5:e19804.

15. Lakhal-Littleton S, Crosby A, Frise M, Mohammad G, Carr C, Loick P, Robbins P. Intracellular iron deficiency in pulmonary arterial smooth muscle cells induces pulmonary arterial hypertension in mice. Proceedings of the National Academy of Sciences. 2019; 116 (26) 13122–13130.

16. Wareing, M. et al. In vivo characterization of renal iron transport in the anaesthetized rat. Journal of Physiology. 2000; 524: 581–586.

17. Veuthey, T. et al. Role of the kidney in iron homeostasis: renal expression of Prohepcdin, Ferroportin, and DMT1 in anemic mice. AJP: Renal Physiology. 2008;295: F1213–F1221.

18. Lakhal-Littleton S, Wolna M, Carr CA, Miller JJ, Christian HC, Ball V, Santos A, Diaz R, Biggs D, Stillion R, Holdship P, Larner F, Tyler DJ, Clarke K, Davies B, Robbins PA. Cardiac ferroportin regulates cellular iron homeostasis and is important for cardiac function. Proceedings of the National Academy of Sciences of the United States of America. 2015;112:3164–3169

